# The expression level of CACNA1C-encoded Ca_V_1.2 is a tipping point between promotion and inhibition of dendritic growth in neurons

**DOI:** 10.64898/2025.12.11.693636

**Authors:** Stefano Lanzetti, Pietro Mesirca, Ninoska Polona Diestra, Eleonora Torre, Martina Mari, Sabrin Haddad, Gerald J. Obermair, Marta Campiglio, Alessandra Folci, Matteo E. Mangoni, Valentina Di Biase

## Abstract

The CACNA1C gene encodes the Ca_V_1.2 L-type voltage-gated calcium channel, which plays a crucial role in neuronal signaling. CACNA1C is a risk gene for psychiatric conditions involving disruption of neuronal connectivity such as schizophrenia, autism, and bipolar disorders. While genomic studies are consistently reinforcing the notion of CACNA1C as an important locus related to these diseases, the role of Ca_V_1.2 channels in determining neuronal architecture is incompletely understood. Several studies pinpoint the L-type current (*I*_CaL_) as regulator of dendritic arborization development. *I*_CaL_ lays upstream of competing cellular mechanisms leading to both inhibition and promotion of dendritic growth. How signal selectivity is achieved remains an open question. Here, we report that *I*_CaL_-dependent dendritic development of murine cultured hippocampal neurons relies on Ca_V_1.2 and is determined by an equilibrium between the level of Ca_V_1.2 protein expression and *I*_CaL_ activity. Indeed, increasing *I_CaL_* enhances dendritic complexity only when Ca_V_1.2 expression level is reduced. In contrast, when channel levels are at baseline, the Ca_V_1.2-dependent growing signal is overcome by the elevation of the dendritic growth inhibiting CaMKIIα signaling beyond basal conditions. These findings suggest that Ca_V_1.2 expression level acts as a molecular switch between dendritic growing and inhibiting signals. Consequently, altered Ca_V_1.2 expression during early development may alter neuronal structure, potentially impairing neural network formation and increasing susceptibility to psychiatric disorders.

**SIGNIFICANCE STATEMENT:** L-type voltage-gated calcium channels (L-VGCCs) allow calcium influx upon membrane depolarization. In neurons, L-VGCCs play a crucial role in regulating gene transcription, synaptic plasticity, and membrane excitability. Our data demonstrate that the L-VGCCs Ca_V_1.2 isoform is a key regulator of early dendritic development. Calcium influx via Ca_V_1.2 is necessary for proper dendritic growth under basal conditions. However, enhanced calcium currents boost dendritic growth only when channel expression levels are reduced. When the amount of Ca_V_1.2 is at baseline, current stimulation antagonizes the growing process by upregulating the CaMKIIα signaling. Our findings suggest that the number of Ca_V_1.2 available determines whether channel activity leads to aberrant dendritic arborization by sizing the recruitment of growth inhibiting CaMKIIα cascade.

## INTRODUCTION

L-type voltage-gated calcium channels (L-VGCCs, Ca_V_1s) provide a major source of calcium entry in excitatory neurons where they control multiple functions, including regulation of gene transcription, synaptic plasticity, and membrane excitability. Calcium currents through Ca_V_1s (*I_CaL_*) were also found to be involved in regulation of dendritogenesis and dendritic arborization (1). This widely accepted channel function still requires elucidation, as results from previous reports lead to unclear conclusions on how the specific underlying pathways are elicited. Indeed, *I_CaL_* has been repeatedly shown to lay upstream of competing mechanisms leading to both promotion and inhibition of dendritic growth within the same neuronal population (2–4).

L-VGCCs stimulate dendritic growth via different signaling pathways including calmodulin kinase IV (CaMKIV), RhoA cascade, and regulation of gene transcription via CREB phosphorylation (2, 4, 5). Intriguingly, L-VGCCs mediate also activity-dependent inhibition of dendritic growth by CaMKIIα signaling (2–4, 6). Upon channel opening, the incoming current generates local calcium microdomains, which are detected by molecular calcium sensors, thereby initiating signaling cascades. In its apo state, the calcium-sensing calmodulin (CaM) directly interacts with the IQ-motif of the VGCCs C-terminus. Following channel opening, calcium binds to CaM, which in turn activates CaMKII tethered to L-VGCCs. This physical interaction ensures CaMKII activation selectively by Ca_V_1s and allows spatiotemporal control of channel dependent signal transduction (7, 8). CaMKIIα is believed to restrict dendritic growth by stabilizing the actin cytoskeleton dynamics or even through an inhibitory crosstalk with CaMKIV (4, 9). Whether dendritic promoting and inhibiting signaling depends on specific Ca_V_1 isoform, or other properties of calcium currents is poorly understood.

Direct comparison of VGCCs mRNA profiles shows that Ca_V_1.2 and Ca_V_1.3 are expressed in the hippocampus of mice from embryonic stage to adulthood. Similarly, cultured hippocampal neurons present with comparable amounts of mRNA for both channels from the plating step in which neurites have still to develop to the full differentiation in which a rich dendritic arborization is established and supports the network activity in culture (10). At protein level, Ca_V_1.2 expression exceeds that of Ca_V_1.3 by tenfold, as measured in adult murine brain (11). Interestingly, Ca_V_1.2 and Ca_V_1.3 were described to carry mutations linked to autism and neurodevelopmental disorders, which are associated with aberrant dendritic arborization (12–15). To date, available data indicate that Ca_V_1.2 may control neuritogenesis in murine neurons at the very initial differentiation stage, before polarization is completed (Kamijo et al., 2018). On the other hand, Ca_V_1.3 controls activity-dependent dendritic remodeling in drosophila and perhaps also in mice (16, 17). Despite both channels are thought to participate in shaping dendritic tree structure, their isoform specific contribution remains unclear.

To study how L-VGCCs modulation affects the development of dendrites at early stages, we employ murine cultured hippocampal neurons, an established model of molecular and cellular mechanisms underlying neuronal functions (18) We manipulated *I_CaL_* by pharmacological, molecular, and gene expression silencing approaches and assessed their impact on dendritic complexity, overall length, and branching capacity. Together, our results show that Ca_V_1.2 is an essential regulator of early dendritic outgrowth in basal conditions. However, increasing channel activity may lead to very different degrees of dendritic complexity and elongation. These differences largely depend on the amount of expressed Ca_V_1.2 channels and likely their ability to recruit CaMKIIα signaling.

## RESULTS

### L-VGCCs agonists and antagonists regulate dendritic growth in cultured hippocampal neurons

To examine the effect of L-VGCCs on dendritic growth, eGFP transfected hippocampal neurons were exposed to dihydropyridine (DHP) channel antagonist nifedipine, DHP agonist Bay-K-8644, or non-DHP agonist FPL 64176, for 48 hours from DIV5 to DIV7 (experimental design, Fig. 1A). We employed Bay-K-8644 and FPL-64176 for their differential agonist effects on Ca_V_1 gating (19). Neurons were then immunolabelled at DIV7 with an antibody directed to the CaMKIIα region containing phosphorylated Thr 286, which identifies the enzyme active form (p-CaMKIIα) (20). This approach was previously used to quantify the level of p-CaMKIIα by L-VGCC in fluorescence imaging experiments (8, 21). In addition, we validated the efficacy of anti-p-CaMKIIα antibodies in HEK293 overexpressing eGFP tagged CaMKIIα and treated with ionomycin to induce CaMKIIα phosphorylation, as previously reported (Fig. S1) (22). After image acquisition we measured dendritic complexity by Sholl analysis, which consists of counting the intersections of dendrites with concentric circles of increasing diameter centered at the soma (Fig. 1B). Because the term dendritic complexity comprises dendritic extension and branching, we also obtained for each individual neuron the total dendritic length and number of branching points as independent values (Fig. 1B).

**Figure 1.**
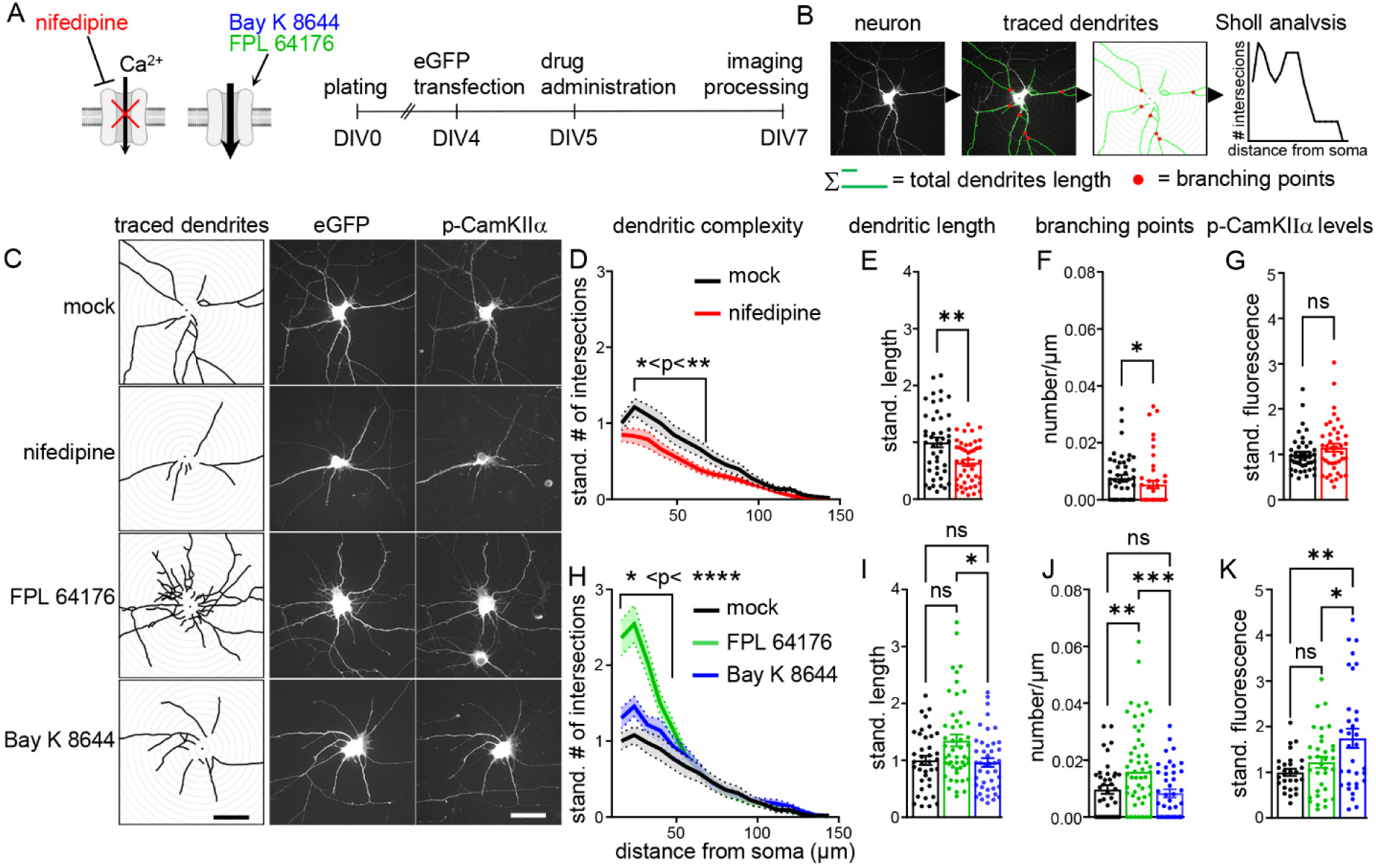
Morphometric analysis of hippocampal neurons treated with 1µM nifedipine, Bay-K-8644, or FPL 64176. **(A)** Experimental design. Hippocampal neurons were transfected with eGFP at DIV4, administered with mock and 1µM nifedipine, 1µM Bay-K-8644, or 1µM FPL-64176 at DIV5, pf fixed at DIV7, and immunolabelled for p-CaMKIIα. (**B**) Flowchart of morphometric measurement process. **(C)** Representative images of eGFP transfected cultured hippocampal neurons, treated with mock, nifedipine, FPL-64176 or Bay-K-8644 with anti-p-CaMKIIα immunolabeling. Bar, 40µm. (**D)** Sholl analysis; nifedipine, red, n=46; mock, black, n=45. Two-Way Anova with Šidák post-hoc test, p-values: Table S1. (**E, F) N**umber of branching points/µm. Mann-Whitney test, **p=0.002, *p=0.03. (**G)** Quantification of p-CaMKIIα immunofluorescence in dendrites; nifedipine, red, n=42; mock, black, n=42. Mann-Whitney test, ns, p=0.226. (**H**) Sholl analysis; mock, black, n =38; FPL 64176, green, n=50; Bay-K-8644, blue, n=43. Two-Way Anova Mixed-model with Tukey post-hoc test, p-values in Table S2. (**I**) Dendritic length; *p=0.024; ns, p>0.999. Kruskall-Wallis followed by Dunn’s post-hoc analysis. (**J**) Branching points/µm; ***p=0.001, **p=0.007; ns, p>0.999. Kruskall-Wallis followed by Dunn’s post-hoc analysis. (**K)** Quantification of p-CaMKIIα immunofluorescence in dendrites; mock, black, n=28; Bay-K-8644, blue, n=33; FPL 64176, green, n=34; **p=0.003, *p=0.034; ns, p=0.580. One-Way ANOVA with Tukey post-hoc test; mean ± SEM.

In line with previous reports (4), nifedipine decreased dendritic complexity, dendritic length, and branching points (Figure 1, red, D, E and F, respectively; statistics, Table S1), confirming the involvement of L-VGCCs in these processes. In the examined neuronal population, p-CaMKIIα levels were similar to mock controls, indicating that CaMKIIα signaling is not required for *I_CaL_* dependent dendritic elongation (Fig. 1G). Unaltered p-CaMKIIα under nifedipine suggests that residual unblocked *I_CaL_* may suffice to maintain basal levels of p-CaMKIIα (Fig. S2).

As nifedipine decreased dendritic growth, the administration of channel agonists Bay-K-8644 and FPL-64176 was expected to increase it. Intriguingly, the dendritic complexity induced by FPL-64176 largely exceeded that recorded with administration of Bay-K-8644 (Fig. 1H, green and blue, respectively; p-values: Table S2). Accordingly, dendritic length and number of branching points upon FPL-64176 treatment were higher than in Bay-K-8644 (Figure 1, I, J). Despite both drugs augmented *I_CaL_* peak, only Bay-K-8644 increased p-CaMKIIα levels, indicating stronger efficiency in recruiting L-VGCCs-dependent CaMKIIα signaling (Fig. 1, K). This result and the observation that Bay-K-8644 induced only a modest increase of dendritic complexity are consistent with the notion that activation of CaMKIIα stabilizes the dendritic arbor and opposes dendritic growth (2–4, 6).

Taken together, these results indicate that activation of *I_CaL_* beyond its baseline level promotes dendritic growth. However, the channel agonists Bay-K-8644 and FPL-64176 enhanced dendritic growth to different extents. Therefore, we investigated whether the underlying reason may depend on differential regulation of *I_CaL_* kinetics beyond the shared effect on current enhancement.

### Bay-K-8644 and FPL-64176 confer distinct kinetics to *I_CaL_* in early cultured hippocampal neurons

To analyze the kinetic properties of *I_CaL_* imparted by Bay-K-8644 or FPL 64176, we measured *I_CaL_* before and upon drugs perfusion (blue and green in Fig. 2, respectively). Representative *I_CaL_* traces are displayed in Fig. 2A and 2F. Averaged current-to-voltage (I-V) relationships showed significant increase of *I*_CaL_ at membrane voltages ranging from −30 mV to + 20 mV for Bay-K-8644 (Fig. 2B, blue) and from −40 mV to −10 mV for FPL-64176 (Fig. 2G, green). Indeed, while both drugs increased *I*_CaL,_ only FPL-64176 significantly shifted current activation to more negative potentials. This result is reflected by the V1/2 activation that was unaltered by Bay-K-8644 (Fig. 2D, blue), but shifted by FPL-64176 (Fig. 2I, green). Mono-exponential fitting of traces from peak to steady state inactivation, recorded at −10, 0, +10 mV, shows that inactivation time constants (τ) were increased by FPL-64176 (Fig. 2H, green), but not by Bay-K-8644 (Fig. 2C, blue), implying that FPL-64176 slows *I_CaL_* inactivation. FPL-64176 and Bay-K-8644 similarly increased *I_CaL_* peak current density normalised to control (Fig. 2E). To evaluate the total calcium entry evoked by 1µM Bay-K-8644 and 1µM FPL-64176 we calculated the current time integral for the peak-corresponding traces. *I_CaL_* time integral measured in FPL-64176 was significantly greater than after Bay-K-8644 perfusion, suggesting a larger time-dependent calcium influx upon FPL-64176 (Fig. 2L). These results align with previous data in which FPL-64176 abolished *I*_CaL_ calcium dependent inactivation (CDI), while Bay-K-8644 increased channel open probability in freshly dissociated neurons from adult mice (19). Since disruption of CDI is a common trait of Ca_V_1 mutations linked to neurodevelopmental disorders characterized by abnormal dendritic arbor (12, 13), we posited that FPL-64176 increased dendritic outgrowth observed in Fig. 1 is determined by slowing *I_CaL_* inactivation rather than by increasing peak current.

**Figure 2.**
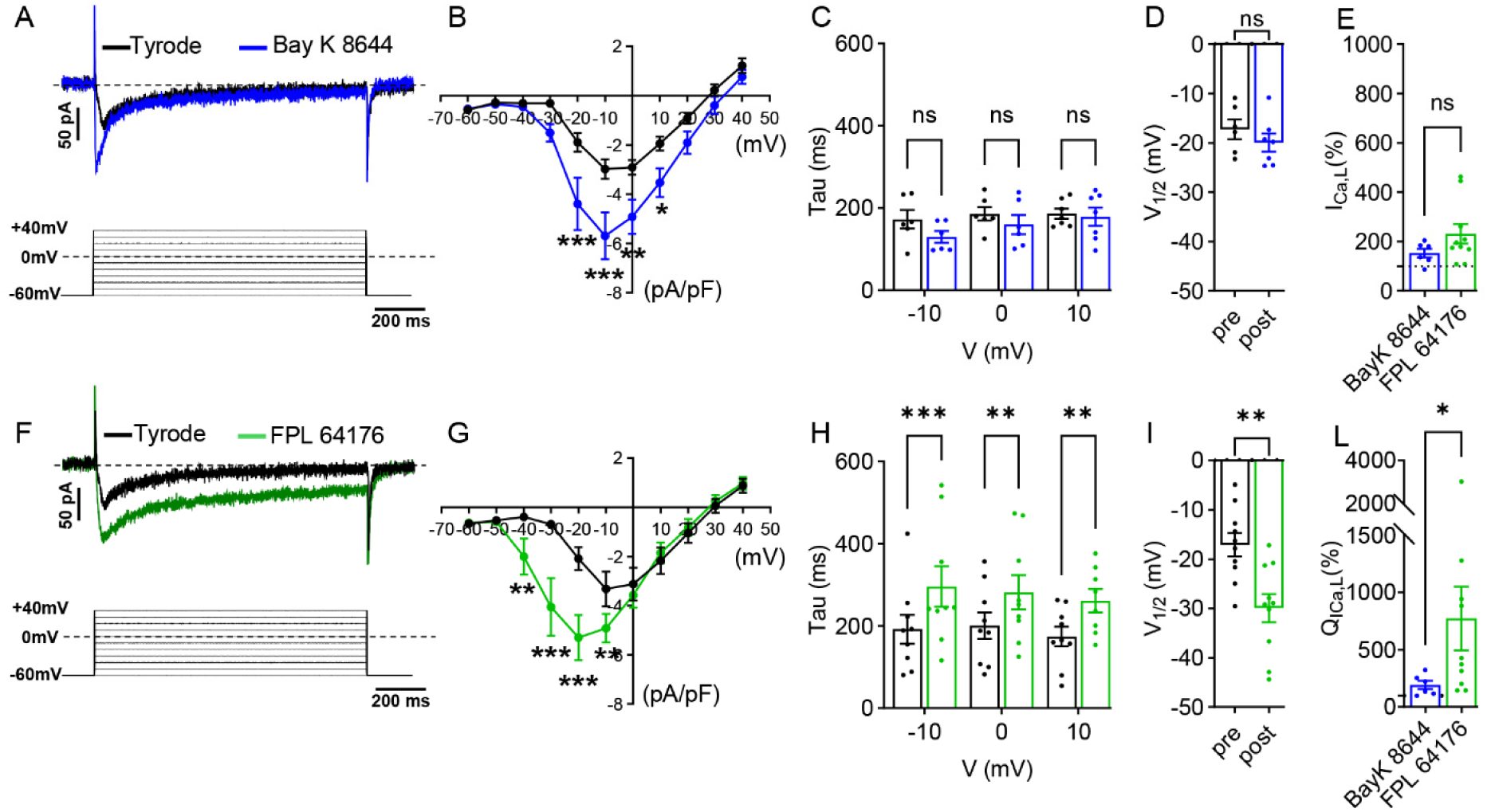
Electrophysiological measurements of *I_CaL_* before and after 1µM Bay-K-8644 or FPL-64176 in young hippocampal neurons. (**A, F**) Representative current traces of *I_CaL_* measured from holding potential −60 mV before (Tyrode, black) and after 1µM Bay-K-8644 (blue) or FPL-64176 (green) perfusion. Voltage-clamp step protocol is reported below. (**B, G**) Current-to-voltage (I–V) relationship of L-type Ca^2+^ current recorded before (*n* = 6/*N* = 4) and after Bay-K-8644 (*n* = 7/*N* = 4) or before and after 1µM FPL-64176 (*n* = 11/*N* = 5). P-values from linear effects mixed model followed by the Tukey’s multiple comparisons test in Table S3. (**C**, **H**) Current decay τ values at −10mV, 0mV and 10mV. Mixed-effects Two-Way ANOVA with Šidák post-hoc test, FPL-64176 *vs* Tyrode at −10mV ***p<0.001 (p=0.004), at 0mV **p=0.05 (p=0.0053), at 10mV **p=0.05 (p=0.0049); Bay-K-8644 *vs* Tyrode at −10mV p=0.220, at 0mV p=0.619, at 10mV p=0.980. (**D, I**) Half-activation voltages (V_1/2_); ns, p=0.3125; **p=0.0020. Wilcoxon test. (**E**) Peak *I_CaL_* (% of control) after 1µM Bay-K-8644 (blue) or 1µM FPL-64176 (green) perfusion. Dotted line 100% control value. Statistics: Mann Whitney test. (**L**) Peak-trace time integral (Q) of *I_CaL_* (% of control) after 1µM Bay-K-8644 (blue) or 1µM FPL-64176 (green) perfusion. Dotted line 100% control value. Statistics: Mann Whitney test; mean ± SEM.

### Dendritic growth is independent of STAC2-induced calcium dependent inactivation

Inhibition of L-VGCC CDI by the Ca_V_1 interactor CaBP1 affects activity dependent neurite outgrowth in cochlear spiral ganglion neurons, suggesting that regulation of channel inactivation may control dendritic development (23). Therefore, we reasoned that CDI disruption may sustain dendritic outgrowth in hippocampal neurons. To test this hypothesis, we took advantage of the ability of Ca_V_1 interacting STAC proteins to abolish CDI (24). Overexpression of the neuronal STAC2 isoform strongly inhibits CDI of endogenous L-VGCC, increasing *I_CaL_* density in cultured hippocampal neurons (25). Using a similar approach (experimental design, Fig. 3A), we overexpressed HA-tagged STAC2 (STAC2-HA) in our model and assessed the influence of CDI inhibition on dendritic growth. To isolate the effects CDI from STAC2 secondary mechanisms, we used the STAC2-ETLAAA-HA construct with a triple alanine substitutions of residues ETL [206-208], which disrupts STAC2-mediated CDI of Ca_V_1.2 and Ca_V_1.3 (26). If reduction of *I_CaL_* CDI promotes dendritic growth, neurons expressing STAC2-HA would grow more than those expressing STAC2-ETLAAA-HA. Any difference with neurons expressing only eGFP would point to STAC2 intrinsic properties independent of CDI. The complexity of neurons expressing the CDI-inactive STAC2-ETLAAA-HA was indistinguishable from that of neurons expressing STAC2-HA (representative neurons in Fig. 3 B; morphometric measurements in Fig. 3 C, D, E, orange and purple, respectively). Together, these results indicate that STAC2-mediated CDI is not a critical factor governing dendritic growth in hippocampal neurons. Despite the abolition of CDI is known to increase *I_CaL_*, CaMKIIα fluorescence levels were unaltered across all conditions, in line with the notion that constraining the recruitment of CaMKIIα signaling generates permissive condition for dendritic growth. These results suggest that slowing *I_CaL_* inactivation kinetics *per se* is dispensable for positive regulation of dendritic outgrowth by FPL 64176.

**Figure 3.**
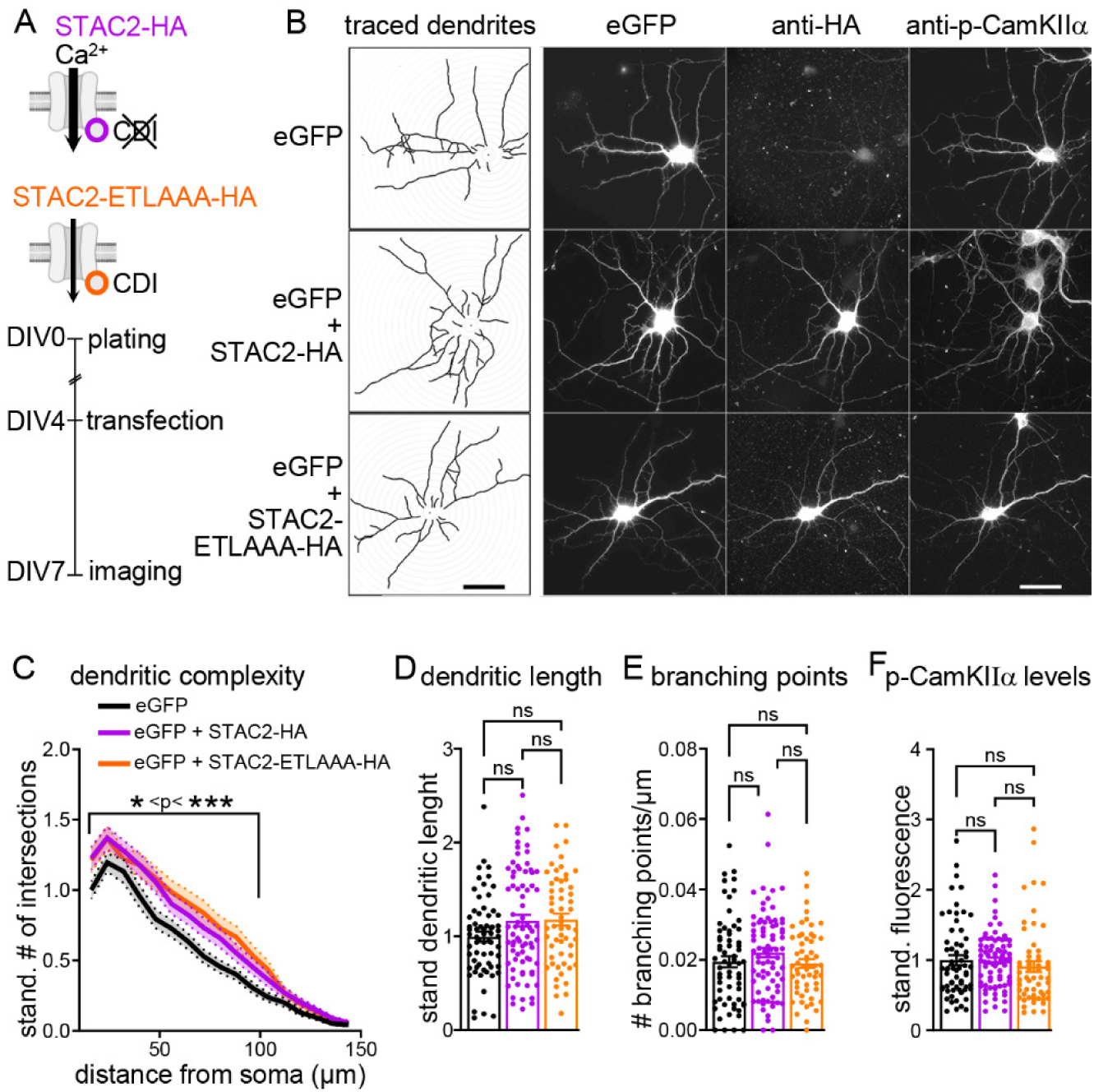
Morphometric analysis of hippocampal neurons transfected with STAC2-HA or STAC2-ETLLAA-HA. (**A**) Experimental design. Hippocampal neurons were transfected at DIV 4 with eGFP plus STAC2-HA or its inactive variant STAC2-ETLAAA-HA. At DIV7 neurons were pf fixed and immunolabeled for HA tag and p-CaMKIIα. **(B)** Representative images of transfected cultured hippocampal neurons showing soluble eGFP fluorescence, anti-HA and anti-p-CaMKIIα immunolabeling. Bar, 40µm. (**C**) Sholl analysis; eGFP, black, n=65; eGFP plus STAC2-HA, purple, n=77; eGFP plus STAC2-ETLAAA-HA, orange, n=57. Two-Way ANOVA Mixed-model with Tukey post-hoc test, p values in Table S4. (**D**, **E**) Dendritic length and branching points/µm. Statistics: **D**, Kruskal-Wallis with Dunn post-hoc test and **E**, ordinary One-Way ANOVA with Tukey post-hoc test. **(G**) Quantification of p-CaMKIIα levels in dendrites. Kruskal-Wallis with Dunn post-hoc test; mean ± SEM.

### Nifedipine and the Ca_V_1.2 selective inhibitor calciseptine similarly reduce dendritic growth

In glutamatergic neurons early dendritic growth depends on the neuronal Ca_V_1.2 isoform, while the Ca_V_1.3 is not involved (27). We thus tested whether inhibition of Ca_V_1.2 using selective antagonist calcispetine aligned with the notion of predominant role of Ca_V_1.2 over Ca_V_1.3 in early dendritic growth. While nifedipine blocks both Ca_V_1.2 and Ca_V_1.3, calciseptine is highly selective for Ca_V_1.2 and spares Ca_V_1.3 (28). If Ca_V_1.2-mediated *I_CaL_* supports dendritic elongation in our system of growing neurons, then calciseptine would restrict dendritic development (Experimental design, Fig. 4A). In addition, any difference between the effect of nifedipine and calciseptine would unmask the potential contribution of Ca_V_1.3 in dendritic growth. Neurons exposed to nifedipine and calciseptine showed diminished complexity in comparison to mock treated ones, with no significant difference between one another (representative neurons Fig. 4B; dendritic complexity Fig. 4C, red and cyan, respectively; p-values: Table S5). This result is reflected by the reduction of dendritic length upon exposure to both drugs leaving the capacity of dendritic branching unaffected (Fig. 4 D and E). These findings confirm that Ca_V_1.2 is the predominant L-type isoform controlling dendritic elongation in our system. This result, together with the lack of evidence in our *I_CaL_* recordings of the typical threshold of Ca_V_1.3-mediated *I_CaL_* in native systems at −40 mV (Fig. 2 B, G, black) (29), led us to check Ca_V_1.3 protein expression. Western blot analysis of Cav1.3 in cultured hippocampal neurons and brain lysates at different developmental stages showed very low detectable Cav1.3 protein at early stages (Fig. S4A). Furthermore, four different sequences of Ca_V_1.3 shRNA failed to diminish the already extremely weak immunostaining against Ca_V_1.3 (Fig. S4C). This observation indicates low or no expression of Ca_V_1.3, corroborating the importance of Ca_V_1.2 channels in L-VGCC-dependent early dendritic development.

**Figure 4.**
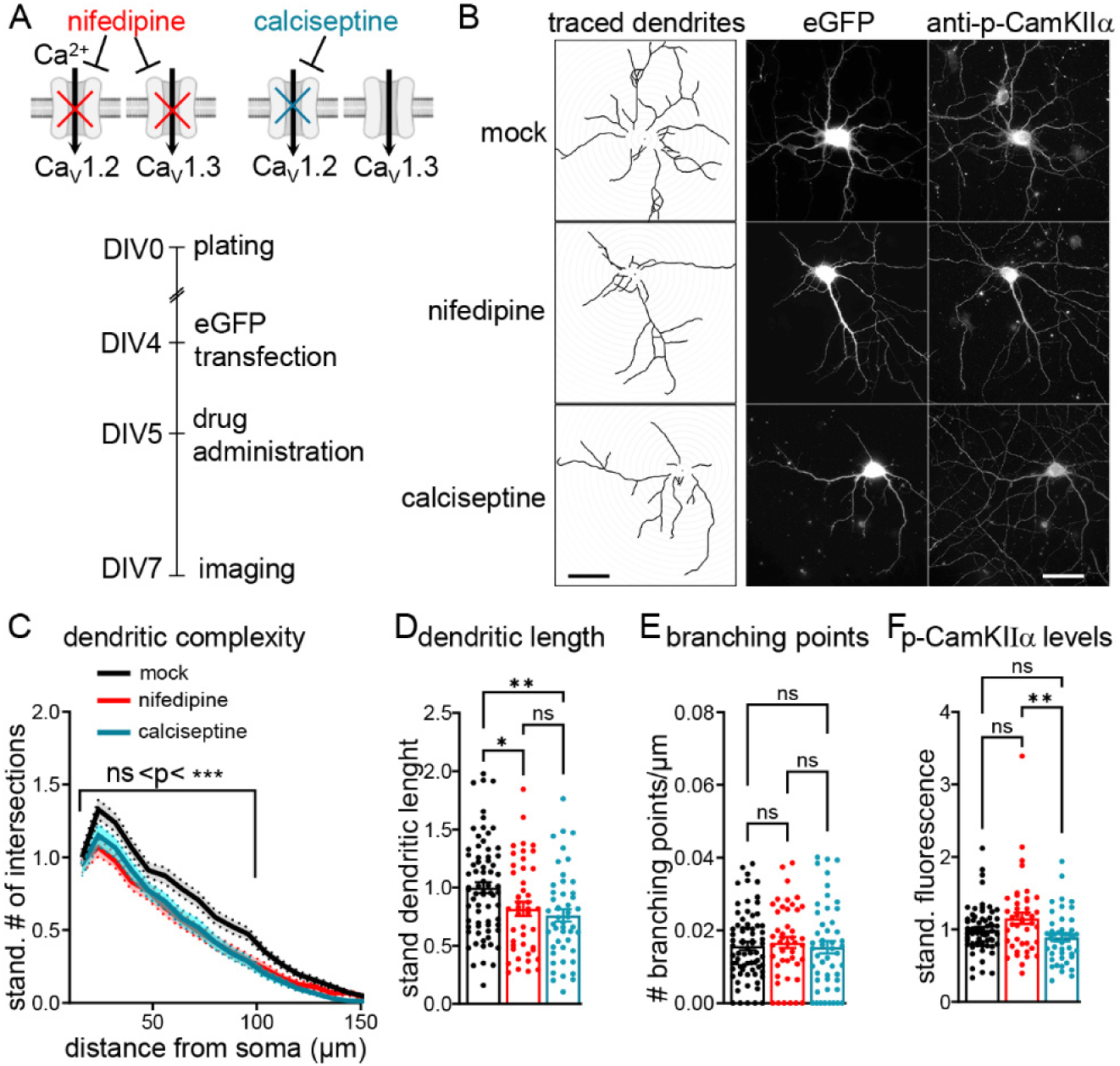
Morphometric analysis of hippocampal neurons treated with 1µM nifedipine or Ca_V_1.2 selective inhibitor calciseptine. (**A**) Experimental design. Hippocampal neurons were transfected with eGFP at DIV4, administered with mock, 1µM nifedipine or 1µM calciseptine at DIV5, pf fixed at DIV7, and immunolabeled for p-CaMKIIα. **(B)** Representative images of cultured hippocampal neurons at the different conditions showing soluble eGFP fluorescence and anti-p-CaMKIIα immunolabeling. Bar, 40µm. (**C**) Sholl analysis; mock, black, n=73; nifedipine, red, n=45; calciseptine, cyan, n=49. Mixed-model Two-Way Anova with Tukey post-hoc test, p values in Table S4. (**D**) Dendritic length. Ordinary One-Way ANOVA with Tukey post-hoc test, *p=0.044; **p=0.004; ns, p=0.774. (**E**) Branching points/µm. Ordinary One-Way ANOVA with Tukey post-hoc test; ns: 0.824 <p < 0.981. (**F**) Quantification of p-CaMKIIα immunofluorescence along dendrites; mock, black, n=66; nifedipine, red, n= 43; calciseptine, cyan, n=46. Kruskal-Wallis with Dunn post-hoc test; **p=0.008; ns: 0.187 < p < 0.446; mean ± SEM.

Quantitative immunofluorescence analysis of p-CaMKIIα is displayed in Fig. 4F. Because CaMKIIα is activated by both Ca_V_1.3 and Ca_V_1.2 (8) immunofluorescence levels of p-CaMKIIα were expected to be higher with calciseptine than with nifedipine, or at most similar, as nifedipine did not reduce p-CaMKIIα levels in our previous experiments shown in Fig. 1. Whereas we confirmed that nifedipine did not reduce p-CaMKIIα levels compared to mock, fluorescence intensity for p-CaMKIIα was surprisingly lower upon calciseptine. These data highlight the efficient inhibition of Ca_V_1.2-dependent CaMKIIα signaling by calciseptine, suggesting that these channels likely exert a prominent role in tuning p-CaMKIIα levels in growing neurons.

### Chronic treatment with FPL-64176 reduces Ca_V_1.2 membrane levels and enhances L-type currents

Administration of Bay-K-8644 and FPL-64176 was shown to internalize Ca_V_1.2 in mature hippocampal neurons (30). To determine whether this is the case also in our experimental settings, we used surface biotinylation assay to measure the Ca_V_1.2 membrane level after 48 hours exposure to 1µM Bay-K-8644, FPL 64176, and nifedipine (Experimental design in Fig. 5A). The immunoblot of total and biotinylated surface fractions of Ca_V_1.2 shows the typical duplet band corresponding to Ca_V_1.2 and its cleaved product (Fig. 5B). To quantify the amount of Ca_V_1.2 in the neuronal membrane, the intensity of biotinylated Cav1.2 was normalized on the transferrin receptor (TfR) signal, which is an independent membrane protein commonly used as reference standard. The absence of the band corresponding to cytosolic calnexin in the biotinylated preparation confirmed the quality of the assay. For all conditions, the values of Ca_V_1.2 at the membrane were expressed as fraction of the internal mock control to compensate for inter-experimental variability (Fig 5C). In presence of FPL-64176, Ca_V_1.2 membrane levels were consistently lower than nifedipine and Bay-K-8644. Next, we tested whether *I*_CaL_ was still increased by FPL-64176 despite the reduced amount of channels at the membrane. Electrophysiology measurements were conducted at the end of the 48 hours treatment and in presence of FPL 64176 or Bay-K-8644 (Experimental design in Fig. 5 A). Representative current waveforms presented in Fig. 5D show that *I_CaL_* peak amplitude upon FPL-64176 and Bay-K-8644 were higher than mock controls and that the effect of FPL-64176 was larger than Bay-K-8644 (Fig. 5D, p-values: Table S6). Thus, despite the lower amount of Ca_V_1.2 channels at the membrane, FPL-64176 still enhanced *I_CaL_* density as Bay-K-8644. Altogether, these results suggest that differences in dendritic complexity induced by Bay-K-8644 and FPL-64176 (Fig. 1) may rely on the reduced level of membrane expressed Ca_V_1.2 channels upon chronic exposure to FPL 64176.

**Figure 5.**
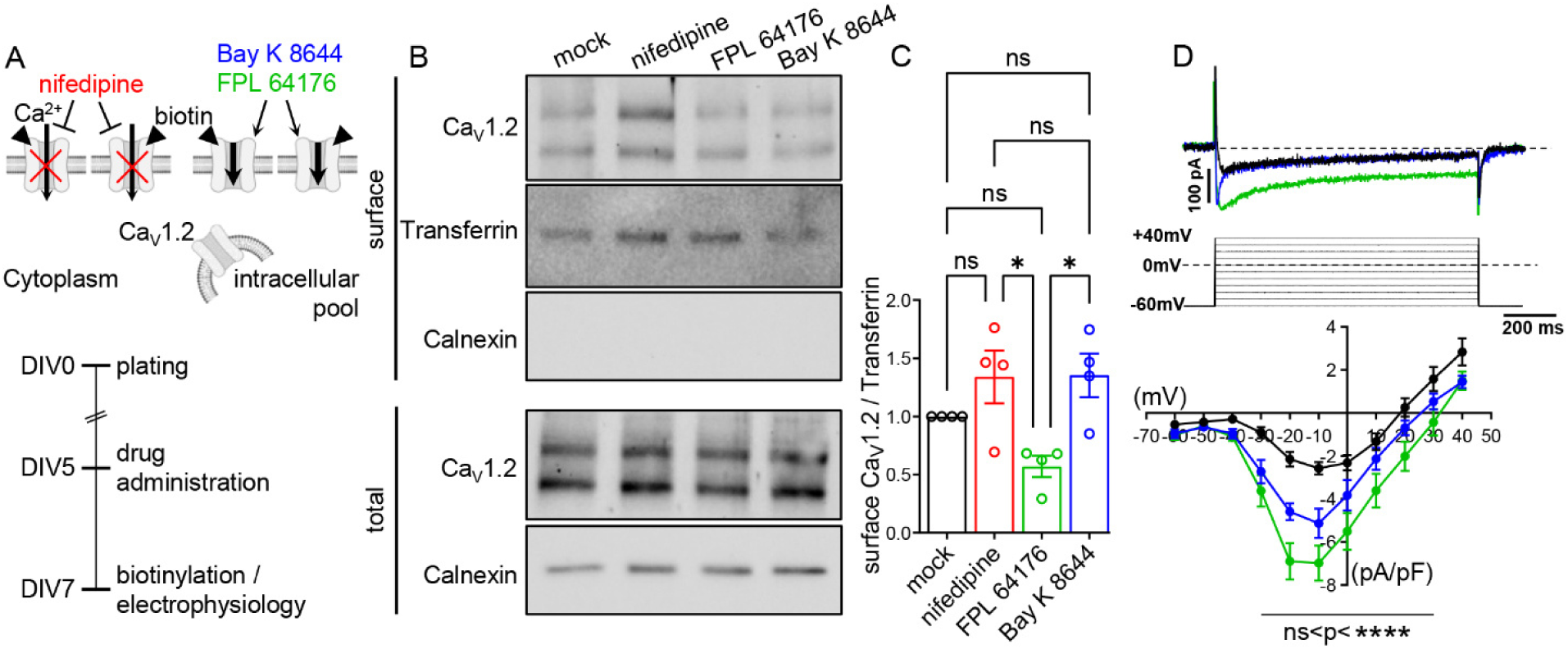
Quantification of Ca_V_1.2 surface expression and *I_CaL_* density after 48 hours administration of 1µM nifedipine, Bay-K-8644, or FPL 64176. (**A**) Experimental design. 1µM nifedipine, 1µM Bay-K-8644, or 1µM FPL-64176 were added at DIV5 into culture dish and maintained for 48 hours. At DIV7 neurons were either processed for biotinylation assay or electrophysiological measurements during perfusion of the corresponding bathing drug. (**B**) Immunoblot showing the surface expressed Ca_V_1.2, TfR loading control, and absence of cytosolic calnexin (upper panel). Lower panel shows the total Ca_V_1.2 in the lysate and calnexin signal as marker of the cytosolic compartment. (**C**) Quantification of Ca_V_1.2 surface signal standardized on TfR intensity and expressed as fraction of mock internal control (black) for neurons treated with FPL-64176 (green), nifedipine (red), and Bay-K-8644 (blue, N=4). Kruskal-Wallis with Dunn post-hoc test, *p=0.043; ns, p>0.999; (**D**) Calcium current traces and voltage-clamp step protocol beneath (upper panel). The lower panel displays I-V curves from holding potential −60mV for mock, black, n=12; Bay-K-8644, blue, n=10; FPL-64176, green, n=12. P-values from linear effects mixed model followed by the Tukey’s multiple comparisons test are given in Table S6; mean ± SEM.

### Reducing Ca_V_1.2 expression level increases dendritic growth upon sustained channel stimulation

To determine how lower Ca_V_1.2 expression level impacts dendritic growth, we knocked Ca_V_1.2 down using specific shRNA. The shRNA efficacy and its effect on the recruitment of CaMKIIα signaling were determined by quantitative immunofluorescence along dendrites. The specificity of the antibody for Ca_V_1.2 versus Ca_V_1.3 was tested in Ca_V_1 null dysgenic GLTs reconstituted with the channel (Fig. S3). Out of four different shRNA sequences weighed, we selected the one yielding Ca_V_1.2 knockdown consistently. Neurons were transfected with pGFP-V-RS-Ca_V_1.2-shRNA plasmid containing the Ca_V_1.2-shRNA and soluble turbo-GFP inserts at DIV4, administered with 1µM Bay-K-8644 at DIV 5, and processed for imaging at DIV7 (Experimental design Fig. 6 A). The extent of Ca_V_1.2 knockdown from scramble to Ca_V_1.2-shRNA is about 30% with cell-to-cell variability (Fig. 6C, scramble + mock black; Ca_V_1.2-shRNA + mock, green; Ca_V_1.2-shRNA + the Bay-K-8644, magenta). First, we observed that neurons exhibiting lower Ca_V_1.2 levels displayed a trend to increase dendritic complexity (Fig. 6B, representative neurons; Fig. 6D, E, F, morphometric measurements, black versus green; p-values for D, Table S7). Second, Ca_V_1.2 knockdown reduced the levels of basal p-CaMKIIα (Fig. 6 G, black versus green), implying that the amount of available Ca_V_1.2 sizes the active pool of CaMKIIα. Third, Bay-K-8644 treatment further enhanced dendritic complexity in neurons expressing Ca_V_1.2-shRNA (Fig. 6 D, E, F, magenta; statistics for D, Table S5) and failed to increase CaMKIIα signaling as we observed in neurons with the Ca_V_1.2 expression levels at baseline (Fig. 1 and 5). These results suggest that the reduction of available Ca_V_1.2 channels prevents wide-scale L-type dependent activation of CaMKIIα signaling upon increased channel activity. Thus, CaMKIIα signaling is then not sufficient to overcome the existing growing signals triggered by enhanced calcium influx through Ca_V_1.2.

**Figure 6.**
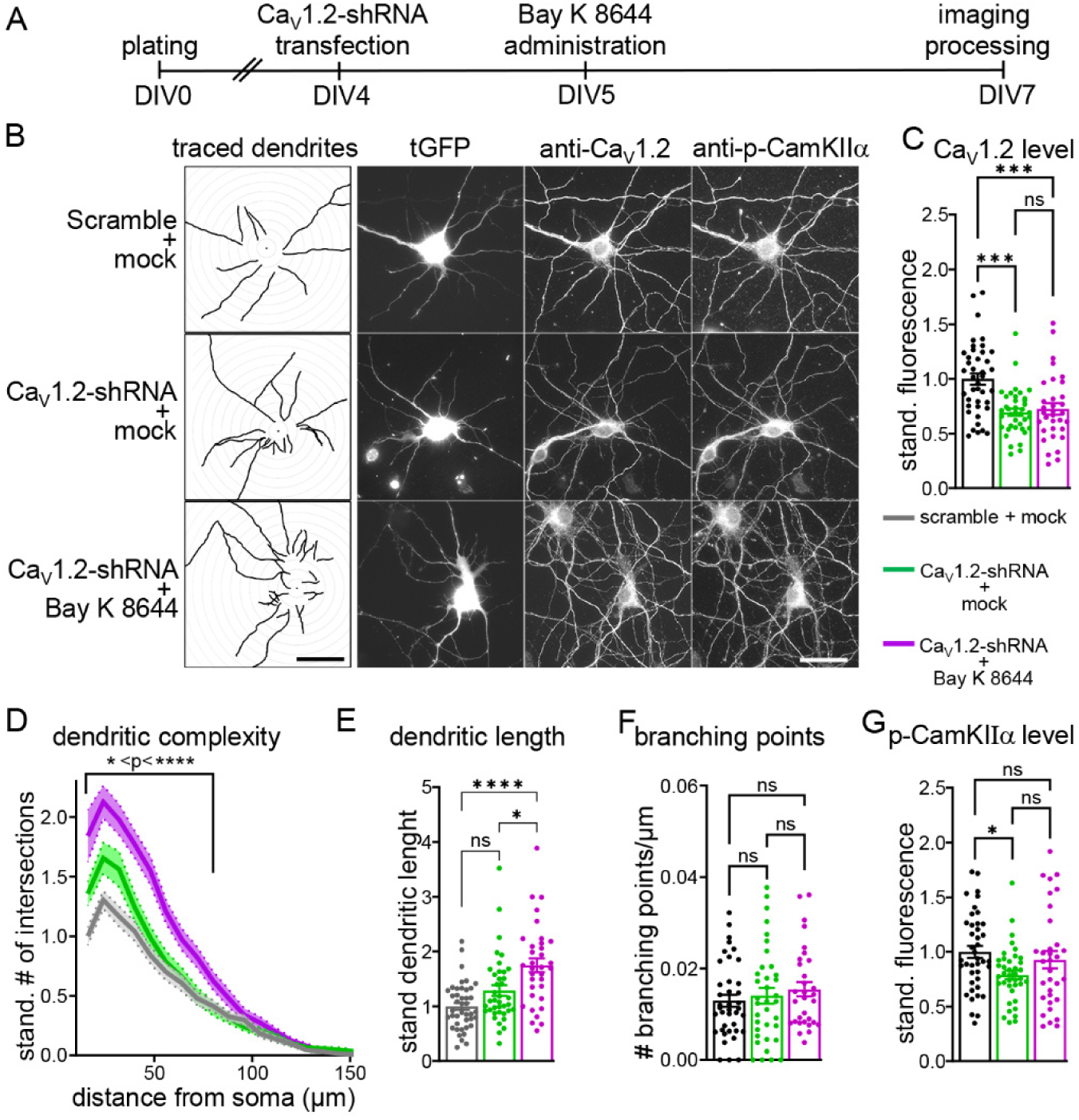
Morphometric analysis of hippocampal neurons transfected with Ca_V_1.2-shRNA and treated with 1µM Bay-K-8644. (**A**) Experimental design. Neurons were transfected scramble-shRNA or Ca_V_1.2-shRNA at DIV4, administered with mock or 1µM Bay-K-8644 at DIV5 for 48 hours, pf fixed at DIV7, and immunolabeled with anti-Ca_V_1.2 and anti-p-CaMKIIα. (**B**) Representative images of cultured hippocampal neurons showing the soluble tGFP fluorescence, with anti-Ca_V_1.2 and anti-p-CaMKIIα immunolabeling. Bar, 40µm. (**C**) Quantification of anti-Ca_V_1.2 immunofluorescence along dendrites; scramble-shRNA plus mock, grey, n=41; Ca_V_1.2-shRNA plus mock, green, n=39; Ca_V_1.2-shRNA plus Bay-K-8644, magenta, n=33; ***p=0.0001; ***p=0.0006. Kruskall-Wallis followed by Dunn’s post-hoc analysis. (**D**) Sholl analysis; scramble-shRNA plus mock, grey, n=42; Ca_V_1.2-shRNA plus mock, green, n=39; Ca_V_1.2-shRNA plus Bay-K-8644, magenta, n=34. Two-Way Anova Mixed-model with Tukey post-hoc test, p-values in Table S7. (**E**) Dendritic length. ****, p<0.0001; *, p=0.0154, ns, p=0.104, Kruskal-Wallis test with a Dunn’s multiple comparison post-hoc test. (**F**) Number of branching points/µm; ns, 0.491<p<0.859. One-Way ANOVA with Tukey post-hoc test. (**G**) Quantification of p-CaMKIIα immunofluorescence along dendrites; *p=0.0311; ns, 0.624 <p<0.715. Kruskall-Wallis followed by Dunn’s post-hoc analysis; mean ± SEM.

## DISCUSSION

Our data highlight a dual role for Ca_V_1.2 channels for regulation of dendritic growth in young glutamatergic neurons. On one side, *I*_CaL_ through Ca_V_1.2 is required for basal dendritic elongation. On the other, enhanced Ca_V_1.2 activity counteracts the elongation by elevating CaMKIIα signaling beyond baseline. However, recruitment of CaMKIIα signaling fails when Ca_V_1.2 expression level is partially depleted, allowing elongation-promoting mechanisms to predominate (Fig. 7). Together, these findings suggest the existence of an equilibrium between Ca_V_1.2 *I*_CaL_ and CaMKIIα activation, which governs the transition between permissive and restrictive states of dendritic development. Disruption of this balance may underlie aberrant neural circuit formation observed in neurodevelopmental disorders associated with CACNA1C dysregulation.

**Figure 7.**
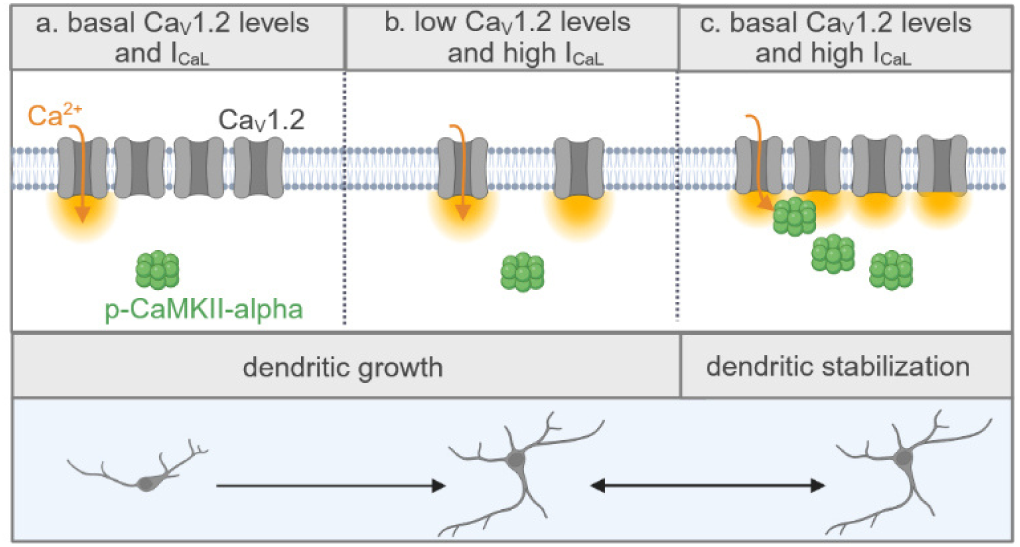
Putative model representing the combined effect of Ca_V_1.2 membrane expression and *I*_CaL_ modulation on dendritic arborization development. (a.) In basal condition calcium entry via Ca_V_1.2 promotes dendritic growth. In this context, the inhibiting CaMKIIα signaling is basal and insufficient to antagonize the development of the dendritic tree. (b.) Increased *I*_CaL_ and low Ca_V_1.2 expression level generate a permissive dendritic growth environment probably by failing further recruitment of CaMKIIα signaling and maintaining the control on promoting signals. (c.) Increased *I*_CaL_ and baseline Ca_V_1.2 level elevate CaMKIIα signaling, counteracting the dendritic growth and stabilizing the dendritic structure.

Propagation of calcium signals and CaMKIIα signaling along dendrites require L-VGCCs, implying that the amount of channel at the membrane is regulated (31, 32). In neurons, Ca_V_1.2 distribute in discrete clusters endowed with a predefined number of slots available for channel occupancy (33, 34). Clustered channels are in a dynamic exchange with a mobile extra-clustered population, which is thought to adjust calcium signaling domains for activity dependent adaptations (33). These channel distribution and dynamics are instrumental to Ca_V_1.2 cooperative channel gating for calcium influx amplification in neurons and cardiomyocytes (35, 36). In this model, calcium influx promotes physical interaction between adjacent channels via Ca2+/CaM calmodulin. Therefore, the overall activity of Ca_V_1.2 within clusters depends on the amount of available channels to form dimers or higher order oligomers. Subtraction of channels from clusters would therefore constrain the cooperative gating and the size of calcium domains. This may explain why enhancing Ca_V_1.2 activity in conditions where the Ca_V_1.2 expression is diminished – as in FPL-64176 treatment and Bay-K-8644 stimulation on shRNA-mediated knockdown – fails to recruit the dendritic growth inhibiting CaMKIIα signaling, resulting in promotion of dendritic growth. Conversely, channel agonist Bay-K-8644 administration on neurons with baseline channel levels increases dendritic-growth inhibiting CaMKIIα signaling.

Our data show that reducing Ca_V_1.2 surface expression increases dendritic complexity. In contrast, cultured hippocampal neurons from heterozygous Cacna1c^+/-^ rats were shown to be less complex than wild type (37). Substantial differences in the experimental model may explain this discrepancy. Cacna1c^+/-^ heterozygote rats present with decreased neurogenesis and conditional Ca_V_1.2 depletion reduces neuronal proliferation rate, causes inefficient differentiation, and hampers cells survival in the hippocampus (38–41). Furthermore, Ca_V_1.2 is essential for proper neuronal migration in brain (27). Because all these Ca_V_1.2 functions may be disrupted in the Cacna1c^+/-^ rats, the resulting neuronal culture maybe enriched with selected neuronal phenotypes (37). Instead, we isolated neurons from wild type mice, preserving the integrity of the neuronal population in culture.

Disruption of CDI characterizes mutations of Ca_V_1.2 and Ca_V_1.3 linked to autism spectrum disorders, which present with aberrant dendritic arborization (12, 13). CDI loss increases I_CaL_, which likely alters cytosolic calcium concentration and potentially modifies calcium signal transduction. Therefore, CDI tuning may adjust calcium dependent functions in excitable cells, including calcium dependent dendritic growth. Consistently, in cochlear spiral ganglion neurons CaBP1 suppression of Ca_V_1 CDI is necessary for excitation-transcription coupling and activity dependent repression of neurites outgrowth (23). However, our study shows that dendritic complexity in hippocampal neurons is unaffected by STAC2-dependent CDI disruption, suggesting that altering CDI does not intrinsically correlate with changes in dendritic complexity per se. It is plausible that dendritic growth modulation by channel CDI might be cell type specific or highly dependent on the signaling microdomain in which channels are embedded.

In our experimental setting, Ca_V_1.3 protein expression is below the threshold of detectability in young neurons. The typical onset of Ca_V_1.3 *I*_CaL_ at −30mV or even −40mV is absent in the I-V curves of mock condition, also observed elsewhere (29, 37). Instead, *I*_CaL_ activation threshold lays at −20mV, the voltage at which Ca_V_1.2 channels open. Ca_V_1.3 very weak immunolabelling intensity (Fig S4 B) did not reduce upon expression of four different Ca_V_1.3 specific shRNA sequences. Furthermore, western blot analysis of brain lysates suggests that Ca_V_1.3 translation could increase as neuronal differentiation progresses (Fig. S4 A). Accordingly, the intensity of fluorescence immunolabelling seems to increase in differentiated neurons (Fig S4 B). This observation complies with the known function of Ca_V_1.3 as fine regulator of membrane excitability in highly specialized systems characterized by well differentiated cell types. Finally, late Ca_V_1.3 expression might concur with the putative role of Ca_V_1.3 for dendritic pruning in murine neurons in response to network adapatation (16, 17, 42).

In conclusion, this study highlights the importance of a tight control of Ca_V_1.2 expression levels in early development of dendrites. Disrupting this critical balance results in aberrant dendritic sprouting and elongation processes that may ultimately alter the foundation of neuronal interconnections.

## MATERIAL AND METHODS

### Low density primary cultures of hippocampal neurons

Low-density cultures of hippocampal neurons were prepared from 16.5-day-old embryonic BALB/c mice either sex as described previously (18, 34). Briefly, dissected hippocampi were dissociated by 2.5% trypsin and trituration. For fluorescence imaging experiments isolated neurons were plated on poly-l-lysine-coated glass coverslips at a density of 3500 cells/cm^2^. After 4 h from plating, coverslips were transferred neuron side-down into a 60-mm culture dish with a glial feeder layer. Neurons and glial feeder layer were cultured in serum-free Neurobasal medium (Invitrogen) supplemented with GlutaMax and B27 (Invitrogen). Neurons were transfected with Lipofectamine 2000 (Invitrogen). For electrophysiology measurements, the same density of neurons from embryonic BALB/c mice was plated on poly-l-lysine-coated 35mm dish (Falcon). For biochemistry experiments low-density cultures of hippocampal neurons were prepared from 18.5-day-old embryonic C57Bl6N mice of either sex. Dissected hippocampi were dissociated in papain (Worthington) for 10 min at 37°C, washed three times and manually triturated in plating medium supplemented with DNAse. Dissociated neurons were plated in 60mm dishes coated with poly-D-ornytine (80 μg/ml, Sigma) at the density of 200.000 cells/dish in MEM supplemented with sodium pyruvate, GlutaMax and penicillin/streptomycin and 10% horse serum. Medium was changed 1 h after plating with Neurobasal supplemented with GlutaMax and B27 (Invitrogen).

### Fluorescence immunolabelling and image acquisition

Neurons were fixed in 4% paraformaldehyde, 4% sucrose in PBS (pF) at room temperature. Fixed neurons were incubated in 5% normal goat serum (NGS) in PBS containing 0.2% bovine serum albumin (BSA) and 0.2% Triton X-100 (PBS/BSA/Triton) for 30 min. Primary antibodies were applied in PBS/BSA/Triton at 4°C overnight or at 4 hours at room temperature. After 3 washes in PBS/BSA/Triton, neurons were incubated for one hours with secondary antibodies to detect fluorochrome-conjugated secondary antibodies (43). Coverslips were mounted in p-phenylenediamine glycerol to retard photobleaching and observed at an Olympus IX73 inverted microscope equipped with a 60X UPLXAPO60XO 1.42 NA oil immersion objective lens, SOLA Light Engine (Lumencor) illumination light source, and Prime BSI back illuminated sCMOS camera (Photometrics). Only images analyzed for data presented in Figure 1 were acquired with an Axio Imager microscope (Carl Zeiss) with a 63× 1.4 NA oil-immersion objective and cooled CCD camera (SPOT; Diagnostic Instruments). To allow quantitative immunofluorescence, images were acquired using the same exposure time and constant illumination intensity within each individual repetition.

### Analysis of fluorescence intensity

Quantification of anti-Ca_V_1.2 and anti-p-CaMKIIα fluorescence intensity was executed using Fiji (https://imagej.net/software/fiji/, (44)). For each transfected neuron, three regions were drawn around dendritic branches using eGFP or tGFP fluorescence as reference, ensuring that neighboring cells were excluded. The regions of interest were then transferred onto the corresponding image of anti-Ca_V_1.2 or anti-p-CaMKIIα immunostaining and the average fluorescence intensity was measured. Background fluorescence intensity was measured for each image. Afterwards, fluorescence values were background subtracted and averaged as a single value for each neuron. These individual values were then standardized on the mean of controls for each repetition. Image acquisition and analysis were blinded, and conditions were unblinded only at the end of the image analysis process.

### Pharmacological treatment and transfection protocols for neuronal morphometric analysis

To analyze the effect of L-VGCCs agonists and antagonists on dendritic growth, cultured hippocampal neurons were transfected at DIV 4 with 0.5µg/dish of pβA-eGFP. Subsequently, they were incubated from DIV 5 to DIV 7 (48 hours) with 1µM FPL 64176 (Sigma Aldrich, cat. # F131-25mg), Bay K 8644 (Sigma, cat. # B112-5m), 1µM nifedipine (Sigma, cat. # N7634), or calciseptine (Alomone, synthetic peptide, cat. # SPC-500) in the culture dish. To study the influence of calcium dependent inactivation of L-VGCCs on dendritic morphology, neurons were transfected at DIV4 with pβA-eGFP (0.5µg/dish), pβA-eGFP (0.5µg/dish) plus pc-STAC2-HA (0.5µg/dish), or pβA-eGFP (0.5µg/dish) plus pc-STAC2-ETLAAA-HA (0.5µg/dish).

To analyze how reduced Ca_V_1.2 expression impacts dendritic growth, neurons were transfected with 1µg/dish with Ca_V_1.2-shRNA constructs (Origene) at DIV 4 and were fixed after 3 days of expression (DIV 7). To examine the consequence of L-VGCCs stimulation in Ca_V_1.2 knockdown neurons, 1µM Bay K 8644 was added to the culture dish from DIV 5 to DIV 7 (48 hours). All constructs were expressed using Lipofectamine 2000 (Invitrogen, cat#11668027) as transfection agent. For all experiment design, neurons were fixed at DIV7 for 10 min at room temperature with pF, and processed for quantitative fluorescence imaging.

### Morphometric measurements

Sholl analysis, dendritic length, and dendritic branching points were conducted using the semiautomated plugin SNT of Fiji (45).

### Antibodies for immunofluorescence

Primary antibodies used in this study were: rabbit anti-Ca_V_1.2 (1:2000, Alomone, cat.# ACC-003, RRID: AB_2039771), guinea-pig anti anti-Ca_V_1.3 (1:1500, Alomone, cat.# ACC-005-GP, RRID: AB_2756614), mouse anti-p-CaMKIIα (1:2000, Invitrogen, cat.# MA1-047, Clone 22B1, RRID: AB_325402, rabbit anti-p-CaMKIIα (1:2000, Cell Signalling, cat.# 12716, clone D21E4F1, RRID: AB_2713889), rat anti-HA (1:500, Roche, cat# 11867423001, RRID: AB_390918). The secondary antibodies (all from Thermo Fischer Scientific) were: goat anti rabbit Alexa 647 (1:4000, Cat# A-21245, RRID:AB_2535813), goat anti guinea-pig Alexa 647 (1:4000, Cat# A-21450, RRID:AB_2535867), goat anti-rabbit Alexa 594 (1:4000, Cat# A-11012, RRID: AB_2534079), goat anti mouse Alexa 594 (1:4000, Cat# A-11032, RRID:AB_2534091), goat-anti rat Alexa 594 (1:4000, Cat# A-11007, RRID:AB_10561522).

### Biotinylation assay

DIV5 neurons were treated with nifedipine (1μM) or FPL 64176 (1μM) or Bay K8644(1μM) for 48h. Controls were treated with vehicle solution. DIV7 treated neurons were first washed twice with warm PBS and then incubated with membrane impermeable EZ-Link Sulfo-NHS-LC-Biotine (0.3 mg/ml in PBS, Thermo Scientific) for 10 min at 4 °C on ice. After three washes with ice-cold PBS, neurons were incubated with 50 mM glycine in PBS for 5 min at 4 °C to quench the remaining unbound biotin. After three more washes in PBS, cells were lysed in lysis buffer (Tris-HCl 10 mM pH 7.5, NaCl 150 mM, EDTA 1 mM, Triton X-100 1%, SDS 0.1%, mammalian protease inhibitor cocktail 1%) for 1 h at 4 °C. After centrifugation (16,000 *g* for 15 min at 4 °C), supernatants containing 150 μg of protein were incubated with 50 μL streptavidin beads overnight at 4 °C to precipitate the surface-biotinylated proteins. After incubation, beads were washed three times with lysis buffer by centrifugation at 5000 × *g* for 5 min at 4 °C. Bead-conjugated membrane proteins were resuspended in sample buffer 2X (350 mM Tris-HCl, pH 6.8, 10% glycerol, 1% SDS, 2.5% β-mercaptoethanol, 25mM DTT, 0.1% bromphenol blue), and loaded onto a 7% acrylamide-bisacrylamide gel. Membranes were then processed as previously described (34). Briefly, membranes were incubated for 1 h with blocking solution (20 mM Tris, pH 7.4, 150 mM NaCl, 0.1% Tween 20, and 5% dried nonfat milk (Regilait) at room temperature and then exposed overnight at 4 °C to the following primary antibodies: rabbit anti-Ca_V_1.2 (1:500, Alomone, Cat. # ACC-3300, RRID: AB_2039771); mouse anti-transferrin (1:1000, Cell Signalling, Cat. BK46222S, RRID: AB_3661976); rabbit anti-calnexin (1:2000, Sigma Aldrich, Cat C4731., RRID: AB_476845). HRP-conjugated secondary antibody (Thermo Scientific) was applied for 1 h in blocking buffer at room temperature. The chemiluminescent signal was developed with Clarity Western ECL Substrate (Biorad) and detected using ChemiDoc^TM^ MP System (Bio-Rad). The intensity of non-saturated band signals was quantified using Fiji software. The levels of cleaved and full-length Ca_V_1.2 were normalized using the intensity of the corresponding Transferrin bands.

### Electrophysiological recordings

Isolated hippocampal neurons were patched in whole-cell configuration with a Molecular Devices 700A amplifier coupled with a Molecular Devices 1550B Digidata. Signals were filtered at 2 kHz via pClamp 10.4 (Molecular Devices). Membrane capacitance (C_m_) and series resistance were measured in all cells. Measurements were performed at physiological temperature (37 °C) with a temperature-controlled chamber and warmed solutions. Calcium currents were recorded with an extracellular solution (pH 7.4) containing (mM): 135 TEA-Cl, 2 CaCl_2_, 1 MgCl_2_, 10 HEPES, 10 4-Aminopyridine, 5.5 glucose, 0.3 tetrodotoxin (TTX). Patch pipettes (resistance 2-3 MΩ) were filled with intracellular solution (pH 7.2) containing (mM): 20 TEA-Cl, 10 HEPES, 125 CsOH, 1.2 CaCl_2_, 5 MgATP, 0.1 LiGTP, 5 EGTA. L-type Ca^2+^ current was activated by applying depolarizing steps to the range of −60/+40 mV from a HP of −60 mV. *I_CaL_* I-V relationships were obtained by measuring peak current during voltage steps. *I_CaL_* inactivation τ decay (−10/0/10 mV) was obtained by fitting *I_CaL_* between current peak (I_0_) and steady-state current (I_SS_) with one phase decay equation:

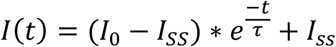

Half-activation voltages (V_1/2_) were calculated by fitting current I-V curves by using a modified Boltzmann equation:

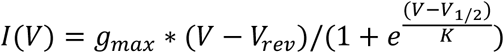

where *V_rev_* is the current reversal potential, V is the membrane voltage, I is the peak current at given voltage, g_max_ is the cell maximum conductance, V1/2 is the voltage for half current activation and K is the slope factor of the Boltzmann relation.

### Plasmids

*pc-STAC2-HA*: To generate STAC2-HA, the coding sequence of mouse STAC2 (Uniprot Q8R1B0) was amplified by PCR using pc-STAC2-GFP (24) as template. The forward primer was designed to introduce a KpnI site, whereas the reverse primer introduced an HA tag and XhoI site. The STAC3-HA fragment generated by PCR was then KpnI/XhoI digested and introduced into the corresponding sites of pc-STAC2-GFP, yielding pc-STAC2-HA.

*pc-STAC2-ETLAAA-HA*: To generate STAC2-ETLAAA-HA, the E206A, T207A and L208A mutations were introduced into STAC2-HA by splicing by overlap extension (SOE) PCR. Briefly, the cDNA sequence of mouse STAC2 (nt 1–1224) was amplified in separate PCR reactions using STAC2-HA as template with overlapping primers mutating the c.617A > G, c.619A > G, c.622C > G and c.623 T > C. The two separate PCR products were then used as templates for a PCR reaction with flanking primers to connect the nucleotide sequences. The resulting fragment was then KpnI/BamHI digested and ligated into the corresponding sites of STAC2-HA.

*p*β*A-eGFP*: previously described soluble eGFP construct (34).

### Ca_V_1.2 shRNA

pGFP-V-RS-Ca_V_1.2-shRNA and scramble control were purchased from OriGene (cat. # TG500242, Locus ID12288 and cat. # TR30013, respectively). The shRNA sequence toward Ca_V_1.2 used in this study reads 5’-TCAGAAGTGCCTCACTGTTCTCGTGACCT-**3’** (Tube ID: TG500242B – GI356982, Origene) while the scramble sequence is 5**’** GCACTACCAGAGCTAACTCAGATAGTACT-3’. Both sequences were cloned into the pGFP-V-RS vector (OriGene, cat. # TR30007) containing turbo-GFP (tGFP) to identify transfected cells.

### Ca_V_1.3 shRNA

Four pGFP-V-RS-Ca_V_1.3-shRNA constructs were purchased from OriGene (cat# TR500243, Locus ID 12289; Ca_V_1.3 shRNA insert were ad hoc cloned into the pGFP-V-RS expression vector, cat # TR30007). The shRNA sequences toward CaV1.3 read as follow: HC110468A: **5’-**TCGTAATCGGCAGCATTATAGACGTGGCC**–3’;** HC110468B: **5’-**TACTTTGACTATGCCTTCACAGCCATCTT**–3’;** HC110468C: **5’-**AGCAGTCCAAGATGTTCAATGATGCCATG**–3’;** HC110468D: **5’-**CTATGCCACTTTCCTGATACAGGACTACT**–3’.**

### Culture of HEK293 cells and p-CaMKIIα activation assay

Cells were grown at 37 °C and 5% CO2 in DMEM plus 10% fetal bovine serum, 1% penicillin/streptomycin, and 1% GlutaMAX (all GIBCO). Cells were co-transfected at 70% confluence with GFP-CaMKIIα using Lipofectamine 2000 (Invitrogen). The GFP-CaMIIα is a gift from Prof. Paul De Koninckv (CERVO Brain Research Center, CA) and is described in (22).

To test the efficacy of our antibodies to recognize the phosphorylated CaMKIIα at Thr-286 we used a previously established protocol. HEK293 cells, transfected or not transfected with GFP-CaMKIIα, were treated with 10µM ionomycin for 2 minutes in culture medium in presence of extracellular calcium. Then fixed immediately fixed in 4% paraformaldehyde and processed for immunofluorescence imaging. For quantitative analysis cytosolic regions were selected based on the GFP fluorescence and the corresponding anti-p-CaMKIIa labelling was measured. Acquisition and analysis were performed in blinded conditions.

### GLTs culture, transfection and immunolabelling

Myotubes from the dysgenic cell line GLT were cultured as previously described (46). Briefly, cells were plated on 13 mm coverslips coated with carbon and gelatine and transfected with 0.5 μg of the desired Ca_V_1 subunit 4 days after plating using FuGENE HD transfection reagent (Promega). GLT myotubes endogenously express the auxiliary α2δ-1, β1a, and γ1 calcium channel subunits as well as STAC3 and the ryanodine receptor, enabling proper functional incorporation of the channel constructs in the triad junction. After 9 days in culture transfected myotubes were fixed with pF and immunolabelled. Primary antibodies used in this study are pan anti-RyR (34-C; 1:250; Alexis Biochemicals, Lausanne, Switzerland), rabbit polyclonal anti-GFP (1:10000, Molecular Probes, Eugene, OR). Cloning procedures for N-terminally GFP-tagged rabbit Ca_V_1.2 (X15539) and human Ca_V_1.3 (EU363339) constructs were previously described (47, 48).

### Western Blot of Ca_V_1.3

Tissue lysates from C75Bl6J were obtained resuspending freshly dissected brains or hippocampi in five volumes (w/v) of ice-cold sucrose buffer (10 mM Tris-HCl pH 7.4, 0.32 M sucrose) supplemented with a protease inhibitor cocktail (Sigma Aldrich, #P8340; 1:1000) and homogenized at 4 °C using a Teflon–glass potter. Nuclear fraction and cell debris were pelleted by centrifugation at 1000 *g* for 10 min. The post nuclear fraction was collected and protein concentration measured using the BCA protein assay (Pierce; #23227). 20μg of protein lysates were loaded onto a 7% acrylamide-bisacrylamide gel and processed as explained for Ca_V_1.2. The primary antibody used to detect Ca_V_1.3 was purchased from Alomone, Cat. #ACC-005, RRID: AB_2039775 and used at 1:500.

### Statistical analysis

Statistical tests were performed using GraphPad Prism version 10.2.2 for Windows (GraphPad Software, San Diego, CA, USA, www.graphpad.com). The number of independent experiments is three to four unless stated otherwise. Data are expressed as mean ± SEM in all figures. Normality was evaluated using D’Agostino & Pearson test. Data set failing the normal distribution were analyzed using non-parametric tests. Statistics are specified in the figure legends.

## Author contribution

S.L., P.M., A.F., E.T., M.M., performed and analyzed experiments. M.C. and G.J.O. shared experimental tools. V.D.B., M.E.M., A.F., and S.L. designed research. V.D.B. conceptualized the study. V.D.B. and S.L. wrote the manuscript, and all other authors edited it.

## ACKNOWLEDGMENTS

We thank Dr. Cornelia Ablinger and Mag. Rosina Maier for help with cell culture and imaging of neurons used for experiments displayed in Figure 1. This work was supported by the Austrian Science Fund (FWF) Grants P332250 (to V.D.B.), P33766 (to M.C.), and by the *Fondation Leducq* TNE 19CV03 to M.E.M. This work is part of the PhD thesis of S.L. Current affiliation of M.M. is Department of Pediatrics at Medical University of Innsbruck.

## SUPPORTING INFORMATION

**Fig S1.**
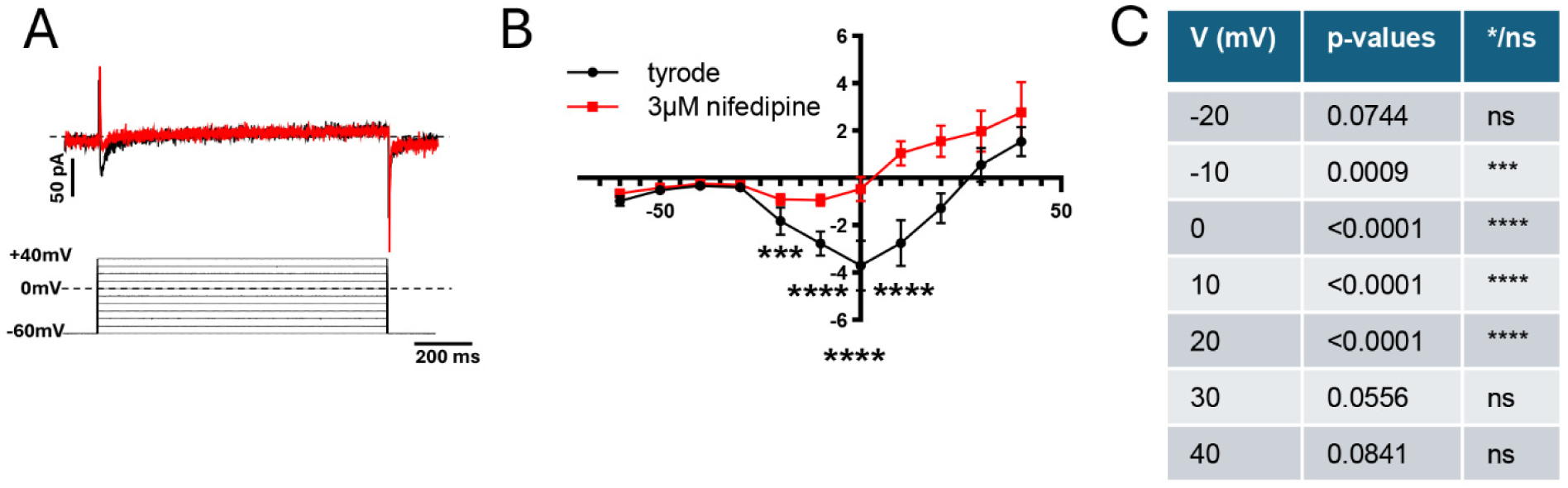
(A). Representative Ca^2+^ current traces and voltage-clamp step protocol. (B) Current-to-voltage (I–V) relationship of L-type Ca^2+^ current recorded from holding potential of −60 mV before (*n* = 6/*N* = 2) and after 3µM Nifedipine (*n* = 7/*N* = 2). (**C**) Statistics: p values from linear effects mixed model followed by the Tukey’s multiple comparisons test.

**Fig. S2.**
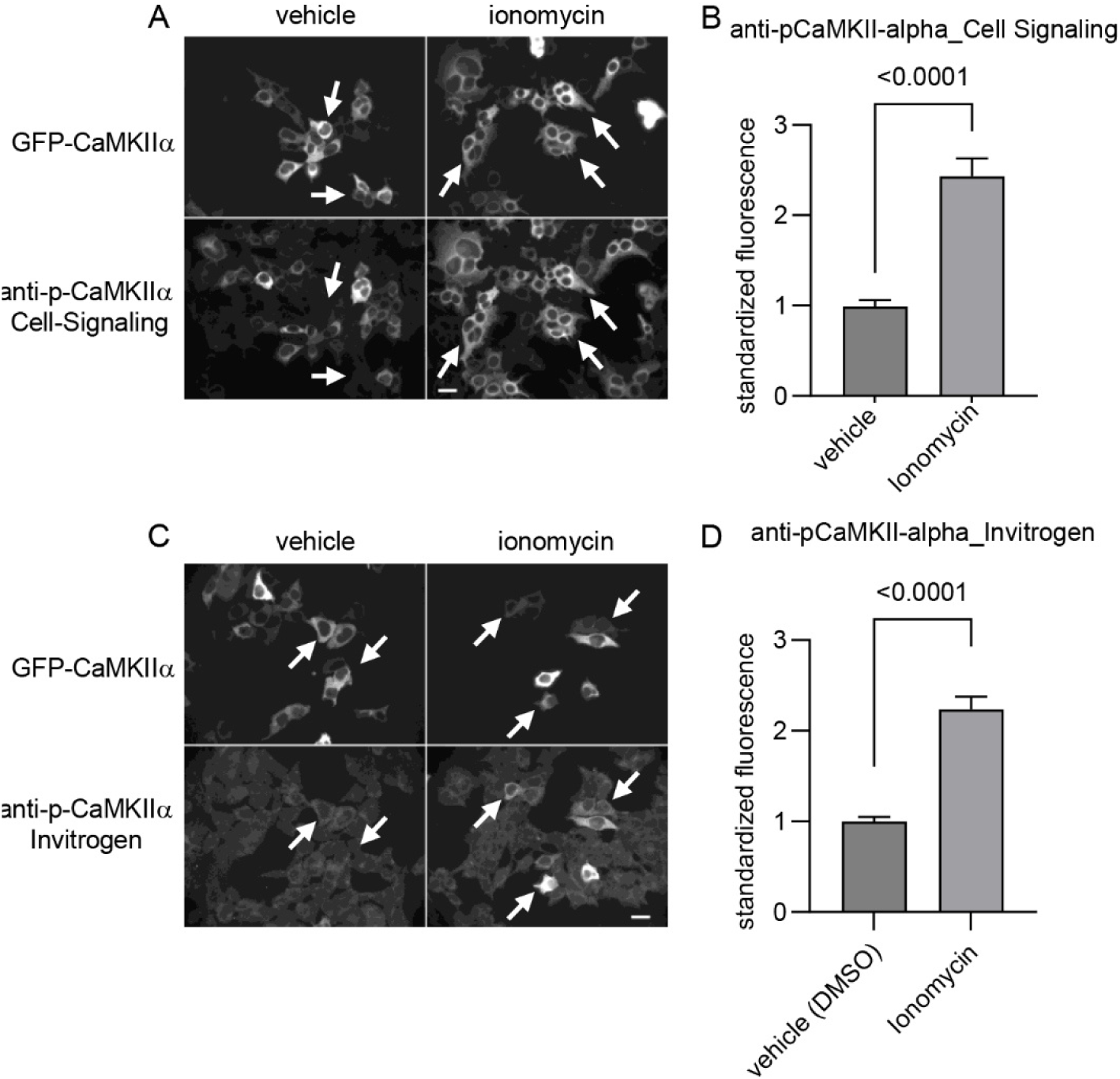
Validation of anti-p-CaMKIIα antibodies used in this study. (A) HEK293 cells expressing eGFP-CaMKIIα treated with vehicle and ionomycin. eGFP-CaMKIIα expressing cells display positive p-CaMKIIα signal (right, arrows) upon ionomycin but not with vehicle (left, arrows). Bar, 20 µm. (B) Bar graphs showing fluorescence quantification of p-CaMKIIα immunolabelling in eGFP-CaMKIIα expressing cells. N=3, Cell Signaling antibody: n_vehicle_= 104, n_ionomycin_= 133, statistics: Mann-Whitney test, p<0.0001; Invitrogen antibody: n_vehicle_= 113, n_ionomycin_= 116 Mann-Whitney test, p<0.0001. Data are presented as mean ± S.E.M.

**Fig. S3.**
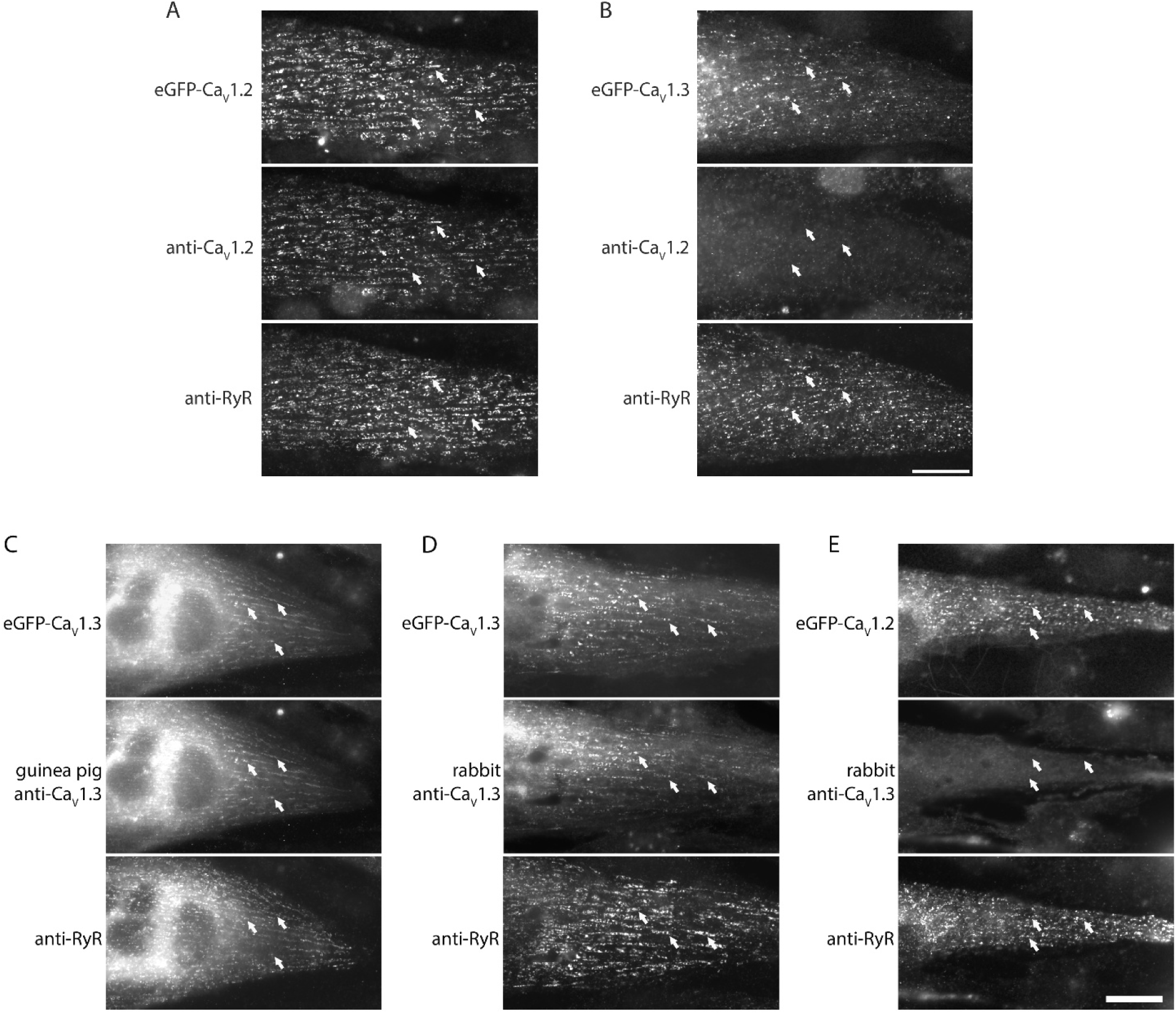
Validation of anti-Ca_V_1.2 and anti-Ca_V_1.3 antibodies used in this study. To test the antibody specificity, we took advantage of dysgenic GLTs skeletal muscle cell line which is null for the Ca_V_1 α pore forming subunit. In this system, heterologously expressed eGFP conjugated Ca_V_1 channels localize at calcium release unites juxtaposed to ryanodine receptors (RyR). Therefore, colocalization between anti-Ca_V_1.2/1.3 and RyR immunolabeling with eGFP fluorescence would indicate the ability of the antibodies to specifically bind to the channel. (A) Anti-Ca_V_1.2 immnunostaning (arrows, middle pannel) corresponds to clusters of heterologous eGFP-Ca_V_1.2 at calcium release units marked by RyR labeling (arrows, lower panel), indicating that the antibody recognizes Ca_V_1.2. (B) Absence of anti-Ca_V_1.2 signal (arrows, middle panel) colocalizing with clusters of eGFP-Ca_V_1.3 (arrows, upper panel) at calcium release units highlighted by RyR labelling (arrows, lower panel) indicates that the antibody does not cross react with the Ca_V_1.3 channel isoform. (C and D) Guinea-pig and rabbit anti-Ca_V_1.3 distribute in clusters (arrows, middle panel) colocalizing with eGFP-Ca_V_1.3 and RyR (arrows, upper and lower panel, respectively), indicating that both antibodies recognize Ca_V_1.3 channels. (E) Absence of anti-Ca_V_1.3 signal (arrows, middle panel) colocalizing with clusters of eGFP-Ca_V_1.2 (arrows, upper panel) at calcium release units highlighted by RyR labelling (arrows, lower panel) indicates that the antibody does not cross react with the Ca_V_1.2 channel isoform. Bar, 20 µm

**Fig. S4.**
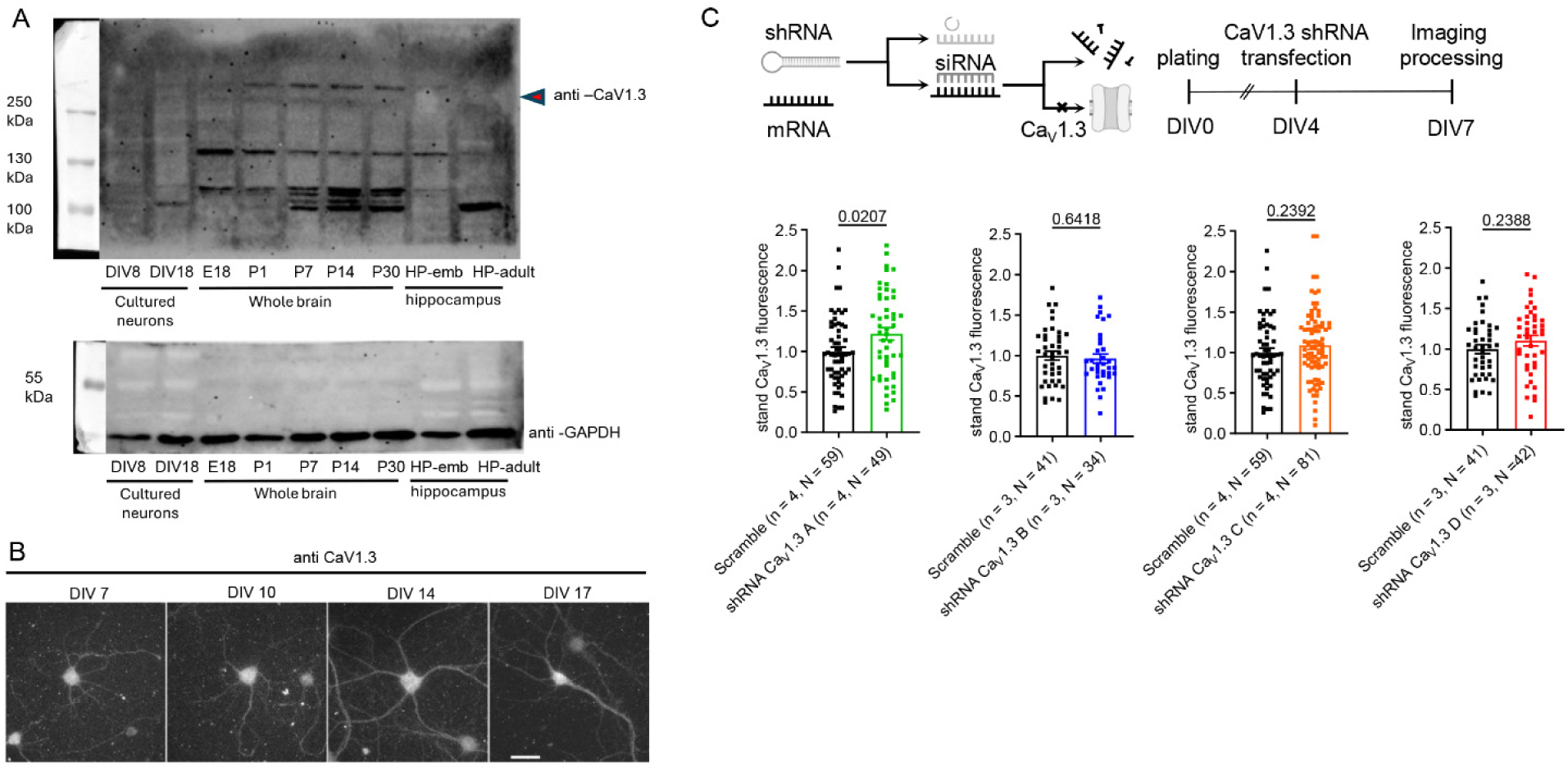
Ca_V_1.3 L-VGCC channel isoform is below detectability with the experimental tools used in this study. **(A)** Immunoblot showing Ca_V_1.3 expression levels in lysates from cultured hippocampal neurons (DIV8 and DIV18) whole brain at embryonic day 18 (E18) and post-natal day (PND) 1, PND7, PND14, PND30 and hippocampus (embryonal and adult). (B) Immunolabeling with anti-Ca_V_1.3 is very weak and difficult to dissect from background at DIV7. At visual inspection, the anti-Ca_V_1.3 immunofluorescence signal seems to progressively increase at the following time-points. Bar, 40 µm. (C) Quantification of anti-Ca_V_1.3 immunofluorescence in DIV7 cultured hippocampal neurons expressing four different shRNA directed to Ca_V_1.3.

**Table S1.** p-values of post-hoc multiple comparison analysis of the Sholl analysis n Fig. 1 D.

| Distance from soma ($\mu\text{m}$) | mock vs nifedipine | |
| --- | --- | --- |
|  | P | */ns |
| 16 | 0.952 | ns |
| 24 | 0.006 | ** |
| 32 | 0.044 | * |
| 40 | 0.024 | * |
| 48 | 0.131 | ns |

Multiple comparisons between neurons transfected with eGFP and treated with mock or nifedipine. P-values were calculated for the number of dendritic intersections with concentric circles of increasing diameters centered on the soma.

**Table S2.** P-values of post-hoc multiple comparison analysis shown in Fig. 1 H.

| Distance<br>from soma<br>($\mu\text{m}$) | Condition | | | | | |
| --- | --- | --- | --- | --- | --- | --- |
|  | mock vs Bay K 8644 |  | mock vs FPL 64176 |  | Bay K 8644 vs FPL 64176 |  |
|  | P | */ns | P | */ns | p | */ns |
| **16** | 0.1926 | ns | <0,0001 | **** | 0.0006 | *** |
| **24** | 0.0851 | ns | <0,0001 | **** | 0.0007 | *** |
| **32** | 0.3979 | ns | 0.0002 | *** | 0.002 | ** |
| **40** | 0.4972 | ns | 0,0172 | ** | 0.2273 | ns |
| **48** | 0.6814 | ns | 0,1016 | ns | 0.4229 | ns |

Multiple comparisons between neurons transfected with eGFP and treated with mock, Bay K 8644 or FPL 64176. P-values were calculated for the number of dendritic intersections with concentric circles of increasing diameters centered on the soma.

**Table S3.** p-values of I-V curves at different voltages before and after Bay K 8644 and FPL 64176 perfusion.

| **I-V curves (N=3)** |  |  |  |  |
| --- | --- | --- | --- | --- |
| **V(mV)** | **Tyrode vs Bay K 8644 (n=7)** |  | **Tyrode vs FPL 64176 (n=9)** |  |
|  | **P** | **\*/ns** | **P** | **\*/ns** |
| **-40** | 0.8295 | Ns | 0.0032 | ** |
| **-30** | 0.0556 | Ns | <0.0001 | *** |
| **-20** | 0.0002 | *** | <0.0001 | *** |
| **-10** | <0.001 | *** | 0.0035 | ** |
| **0** | 0.0017 | ** | 0.4045 | ns |
| **+10** | 0.0133 | * | 0.553 | ns |

Multiple comparisons were conducted between $I_{CaL}$ elicited at increasing voltage steps on neurons treated with mock, 1 μM Bay K 8644, or 1 μM FPL 64176.

**Table S4.** P-values of post-hoc multiple comparison analysis shown in Fig. 3C.

| Distance<br>from soma<br>($\mu\text{m}$) | eGFP vs eGFP +<br>STAC2-HA | | eGFP vs eGFP +<br>STAC2-ETLAAA-HA | | eGFP +STAC2-HA vs<br>eGFP + STAC2-ETLAAA-HA | |
| --- | --- | --- | --- | --- | --- | --- |
|  | P | */ns | P | */ns | p | */ns |
| 16 | 0.0107 | * | 0.0142 | * | 0.9949 | ns |
| 24 | 0.0623 | ns | 0.0486 | * | 0.9600 | ns |
| 32 | 0.1358 | ns | 0.4767 | ns | 0.8003 | ns |
| 40 | 0.0092 | ** | 0.0143 | * | 0.9974 | ns |
| 48 | 0.0011 | ** | 0.0010 | ** | 0.9551 | ns |
| 56 | 0.0381 | * | 0.0023 | ** | 0.5008 | ns |
| 64 | 0.0191 | * | 0.0012 | ** | 0.5408 | ns |
| 72 | 0.0113 | * | 0.0004 | *** | 0.4399 | ns |
| 80 | 0.0130 | * | 0.0018 | ** | 0.6842 | ns |
| 88 | 0.0678 | ns | 0.0025 | ** | 0.3846 | ns |
| 96 | 0.0784 | ns | 0.0090 | ** | 0.5897 | ns |
| 104 | 0.1793 | ns | 0.0397 | * | 0.6904 | ns |
| 112 | 0.5502 | ns | 0.6293 | Ns | 0.9985 | ns |

Multiple comparisons between neurons transfected with eGFP, eGFP plus STAC2-HA, or eGFP plus STAC2-ETLAAA-HA. P values were calculated for the number of dendritic intersections with concentric circles of increasing diameters centered on the soma.

**Table S5.** P-values of post-hoc multiple comparison for Sholl analysis shown in Fig. 4C.

| Distance from soma (μm) | mock vs nifedipine |  | mock vs calcisepetine |  | nifedipine vs calcisepetine |  |
| --- | --- | --- | --- | --- | --- | --- |
|  | P | */ns | P | */ns | p | */ns |
| 16 | 0.9294 | ns | 0.7776 | ns | 0.9595 | ns |
| 24 | 0.0001 | *** | 0.0107 | * | 0.4421 | ns |
| 32 | <0.0001 | *** | 0.0125 | * | 0.3845 | ns |
| 40 | 0.0005 | *** | 0.0307 | * | 0.4636 | ns |
| 48 | 0.0564 | ns | 0.0655 | ns | 0.9935 | ns |
| 56 | 0.0011 | ** | 0.0120 | * | 0.7628 | ns |
| 64 | 0.0014 | ** | 0.0029 | ** | 0.9627 | ns |
| 72 | 0.0013 | ** | 0.0048 | ** | 0.9096 | ns |
| 80 | 0.0164 | * | 0.0135 | * | >0.9999 | ns |
| 88 | 0.0165 | * | 0.0065 | ** | 0.9767 | ns |
| 96 | 0.0104 | * | 0.0052 | ** | 0.9913 | ns |
| 104 | 0.1680 | ns | 0.0338 | * | 0.8336 | ns |
| 112 | 0.3008 | ns | 0.1320 | ns | 0.9300 | ns |

Multiple comparisons between neurons treated with mock, nifedipine, or calcisepetine. P values were calculated for the number of dendritic intersections with concentric circles of increasing diameters centered on the soma.

**Table S6.** P-values of post-hoc multiple comparison analysis shown in Fig. 5D.

| V (mV) | Tyrode vs Bay K 8644 |  | Tyrode vs FPL 64176 |  | Bay K 8644 vs FPL 64176 |  |
| --- | --- | --- | --- | --- | --- | --- |
|  | P | */ns | P | */ns | P | */ns |
| **-30** | 0.0249 | * | 0.0001 | *** | 0.3638 | ns |
| **-20** | 0.001 | ** | <0.0001 | **** | 0.0010 | ** |
| **-10** | 0.0002 | *** | <0.0001 | **** | 0.0116 | * |
| **0** | 0.0356 | * | <0.0001 | **** | 0.0255 | * |
| **10** | 0.3689 | ns | 0.0005 | *** | 0.0569 | ns |
| **20** | 0.2437 | ns | 0.0005 | *** | 0.1061 | ns |
| **30** | 0.2091 | ns | 0.0047 | ** | 0.3210 | ns |

Multiple comparisons were conducted between $I_{CaL}$ elicited at increasing voltage steps on neurons after 48 hours with mock, 1 μM Bay K 8644, or 1 μM FPL 64176 during perfusion of the corresponding treatment drug.

**Table S7.** P-values of post-hoc multiple comparison analysis shown in Fig. 6 D.

| Distance from<br>soma ($\mu\text{m}$) | A. Sholl analysis | | | | | |
| --- | --- | --- | --- | --- | --- | --- |
|  | shRNA scramble +mock<br>vs<br>Cav1.2 shRNA + mock |  | shRNA scramble +mock<br>vs<br>Cav1.2 shRNA +<br>Bay K 8644 |  | Cav1.2 shRNA + mock<br>vs<br>Cav1.2 shRNA +<br>Bay K 8644 |  |
|  | P | */ns | P | */ns | p | */ns |
| **16** | 0.0002 | *** | <0.0001 | **** | <0.0001 | **** |
| **24** | 0.0003 | *** | <0.0001 | **** | <0.0001 | **** |
| **32** | <0.0001 | **** | <0.0001 | **** | <0.0001 | **** |
| **40** | 0.0617 | ns | <0.0001 | **** | <0.0001 | **** |
| **48** | 0.2240 | ns | <0.0001 | **** | <0.0001 | **** |
| **56** | 0.5996 | ns | <0.0001 | **** | <0.0001 | **** |
| **64** | 0.8424 | ns | 0.0002 | *** | 0.0024 | ** |
| **72** | 0.9452 | ns | 0.0006 | *** | 0.0022 | ** |
| **80** | 0.9679 | ns | 0.0571 | ns | 0.0348 | * |
| **88** | 0.9868 | ns | 0.2299 | ns | 0.1842 | ns |

Multiple comparisons between neurons transfected and treated as follows: scramble shRNA plus mock; Cav1.2 shRNA plus mock, and Cav1.2 shRNA plus 1 $\mu\text{M}$ Bay K 8644. P values were calculated for the number of dendritic intersections with concentric circles of increasing diameters centered on the soma.

## Notes

### Competing Interest Statement

The authors have declared no competing interest.

